# Structural Insights and Inhibitor Discovery for Kyasanur Forest Disease Virus NS5 Methyltransferase

**DOI:** 10.64898/2026.08.14.744817

**Authors:** Pratibha Verma, Arpan Kayastha, Preeti Dhaka, Mandar Bhutkar, Pravindra Kumar, Shailly Tomar

## Abstract

Kyasanur Forest Disease Virus (KFDV) NS5 methyltransferase (MTase) protein is the essential enzyme that is involved in the cap methylation of viral RNA, viral replication, and immune evasion, and therefore it is an important protein of interest for antiviral research and drug design. In the present work, we successfully resolved the three-dimensional crystal structures of KFDV NS5 MTase co-crystallised with SAH and GTP at resolutions of 2.2 Å and 2.6 Å, respectively. In previous studies, HC (Herbacetin) and CAPE (Caffeic acid phenethyl ester) have shown inhibitory activity against SAM-dependent viral MTase. To evaluate the inhibitory potential of HC and CAPE against KFDV NS5 MTase, we have performed isothermal titration calorimetry (ITC) and tryptophan fluorescence spectroscopy (TFS) to validate protein interaction with target compounds. MTase inhibition assay was performed using capillary electrophoresis (CE) assays. Additionally, fluorescence polarisation (FP) confirmed RNA binding inhibition by CAPE and HC. Together, these experiments suggest that HC and CAPE are promising inhibitors against KFDV NS5 MTase and could potentially act as lead compounds to design broad-spectrum anti-Orthoflavivirus drugs.

## 1. Introduction

*Orthoflavivirus kyasanurense*, also known as Kyasanur Forest disease virus or monkey fever, belongs to the *Orthoflavivirus genus* (formerly known as Flavivirus) in the family *Flaviviridae*, and causes Kyasanur Forest disease (KFD). This genus encompasses major human viral pathogens, Orthoflavivirus zikaense (ZIKV), Orthoflavivirus dengue serotypes 1-4 (DENV 1-4), Orthoflavivirus japonicum (JEV), Orthoflavivirus encephalitidis (TBEV), Orthoflavivirus flavi (YFV), and Orthoflavivirus nilense (WNV) (Simmonds et al., 2017). KFDV belongs to a closely related group of viruses that includes Nanjianyin virus and Alkhurma hemorrhagic fever virus (AHFV) (Thomas et al., 2014). In 1957, in the Kyasanur Forest region of the Indian state, Karnataka, the KFD virus was isolated from a diseased monkey (Work et al., 1959). It is sustained in enzootic cycles that include monkeys, rodents, and shrews as amplifying hosts (Work et al., 1959).

Among the nonhuman primates (NHPs), KFDV causes epizootic outbreaks that lead to high fatality, particularly in *Semnopithecus entellus* (formerly known as *Presbytis entellus*) and *Macaca radiata* (Chakraborty et al., 2019; Work et al., 1959). Between 1957 and 2017, over 3,300 deaths of monkeys were documented in the endemic regions of KFDV in the Western Ghats, India (Sunagar et al., 2026). Notably, from 1964 to 1973, about 1046 cases were recorded (860 *S. entellus* and 186 *M. radiata*) (Sreenivasan et al., 1986; Sunagar et al., 2026). The *Hemaphysalis spinigera* acts as the primary carrier for the transmission of KFDV (Chakraborty et al., 2019; Gupta et al., 2021). Till now, for KFDV, interhuman infection has not been reported (Balasubramanian et al., 2021). Human infections peak seasonally from November to June, underscoring their public health threat in endemic areas (Gupta et al., 2021).

Recently, KFDV has expanded beyond its established endemic area reach in the Western Ghats. Bandipur National Park of Karnataka documented a total of twelve monkeys died in November 2012, and subsequently, KFD symptoms were observed in six human contacts who handled the monkey carcasses (Mourya et al., 2013). Between November 2012 and May 2013, KFDV infection was confirmed in human samples tested from the Maddur Forest Range in Karnataka. Later, there were outbreaks in Kerala (Wayanad in 2013 and Malappuram in 2014), Tamil Nadu (Nilgiri in 2013), Goa (Sattari taluk in 2015), and Maharashtra (Sindhudurg district in 2016) (Mourya et al., 2013; Tandale et al., 2015).

The KFD is a biphasic disease in humans. The first phase includes symptoms including headache, fever, hemorrhagic symptoms, and subsequently second phase is followed by neurological symptoms such as meningoencephalitis (Gupta et al., 2021, 2022). The Orthoflavivirus replication process is complete in the cytoplasm of the target cell. After receptor-mediated, clathrin-dependent internalization, the acidic milieu of the endosome induces structural rearrangement in the envelope (E) glycoprotein, which facilitates fusion of virus and endosome membranes and subsequent liberation of KFDV RNA into the cytoplasm (Smit et al., 2011). Reportedly, mortality rates for KFDV infection lie between 2% and 10% (Munivenkatappa et al., 2018). A KFDV vaccine has been designed from tissue culture and formalin-inactivated, and since 1990, has been in use for several years in the endemic regions of Karnataka, India (Kasabi et al., 2013). The Vaccination has been administered in the outbreak-prone zones; it is given in two doses as a primary vaccination and then boosters at 6 to 9 months intervals (Kasabi et al., 2013). However, it has been proven that the efficacy of this vaccine is limited, and sustained protection is achieved through booster doses (Srikanth et al., 2023). Earlier, it has been reported that the efficacy of the vaccine in a single dose is 62.4% and in two doses is 82.9% (Dandawate et al., 1980, 1994; Kasabi et al., 2013). Currently available Vaccines for KFDV do not provide complete protection, and there are no approved antiviral treatments to prevent or cure KFDV infection (Kasabi et al., 2013). ICMR-National Institute of Virology (NIV), India, and Indian Immunologicals Limited (IIL) have developed a novel whole-virion, attenuated, and adjuvanted vaccine candidate for Kyasanur Forest Disease Virus (KFDV), which shows protective immunity in mice and other animal models and has now entered Phase I clinical trials to investigate safety and immunogenicity in humans with two doses given at an interval of 28 days (Sunagar et al., 2026).

KFDV has an icosahedral nucleocapsid and is a spherical virus measuring 40-65 nm in diameter. The genome of KFDV has a plus-strand, single-stranded RNA genome with 10,774 nucleotides and a large open reading frame (ORF) surrounded by 5′ and 3′ non-translated regions (NTRs) that normally contain structured elements, which regulate replication and protein synthesis (Dodd et al., 2011). The KFDV genomic RNA translates into a single protein chain of 3416 amino acids, that proteolytically process and translate into three structural proteins-capsid (C), premembrane (prM), and envelope (E), and seven non-structural proteins NS1, NS2A, NS2B, NS3, NS4A, NS4B, and NS5 (Dodd et al., 2011). Among Orthoflavivirus proteins, the most conserved and largest non-structural protein is NS5 with a molecular mass of approximately 105 kDa. (da Fonseca et al., 2017; Goh et al., 2024). In recent times, structural studies have revealed as full-length NS5 protein from mosquito-borne Orthoflaviviruses revealed NS5 exists as a dimer; For example, Dengue Virus non-structural protein 5 (DENV NS5, PDB ID 5CCV) (Klema et al., 2016), ZIKA virus non-structural protein 5 (ZIKV NS5, PDB ID 6I7P) (Ferrero et al., 2019), Japanese encephalitis virus non-structural protein 5 (JEV NS5, PDB ID 4K6M) (Lu & Gong, 2013), Yellow Fever Virus non-structural protein 5 (YFV NS5, PDB ID 7QSA) (Dubankova & Boura, 2019), Tick-borne Encephalitis Virus non-structural protein 5 (TBEV NS5 MTase, PDB 7D6M) (Yang et al., 2021), Omsk hemorrhagic fever Virus non-structural protein 5 (OHFV NS5 MTase, PDB 7FJT) (Yang et al., 2021), Langat virus non-structural protein 5 (LGTV NS5 MTase, PDB 7WNJ) (Li et al., 2022). The dimerization of NS5 contributes to NS5 protein stabilization and coordinates RNA capping and RNA synthesis (Klema et al., 2016). This bifunctional NS5 protein consists of two domains, methyltransferase (MTase) at the N-terminal and RNA-dependent RNA polymerase (RdRp) at the C-terminal, which are joined with an adjustable linker of 10 amino acids, in the form of a 3_10_ helix (Ferrero et al., 2019; Flory et al., 2021; Zhao, Soh, Chan, et al., 2015).

The functional RdRp domain is responsible for the *de novo*, template-dependent phosphodiester bond formation in the presence of divalent metal ions and nucleoside triphosphates (NTPs). The functional domain has a typical structure resembling a right hand surrounding the active site, divided into three topologically distinct subdomains: thumb, palm, and fingers. The positively charged residues forming the access pockets are involved in the entry +of NTP substrates (GTP) and template RNA into the catalytic site (Lu & Gong, 2017; Wu et al., 2015; Yap et al., 2007). Inefficiency in the proofreading activity of orthoflavivirus RdRp may lead to the introduction of errors in the viral genome. The significance of NS5 RdRp in genome replication renders it a potential antiviral target (Elena & Sanjuán, 2005; Fernandes et al., 2021; Venkataraman et al., 2018).

The N-terminal NS5 methyltransferase (MTase) domain of Orthoflaviviruses is a SAM-dependent viral MTase. It has three highly conserved substrate-binding pockets: the SAM-binding site, GTP/cap-binding site, and RNA-binding pocket (Egloff et al., 2002a; Zhou et al., 2007). In the SAM-binding site, the evolutionarily stable K-D-K-E (Lys-Asp-Lys-Glu) motif and surrounding residues position S-adenosyl-L-methionine (SAM), which is a universal methyl donor, so that its methyl group can be transferred efficiently to the viral RNA cap. MTase-mediated RNA modification contributes protection and stability of the viral RNA, which facilitates its translation and circumvents innate immunity. (Egloff et al., 2002a; Zhou et al., 2007). The GTP/cap-binding pocket, which exists at N-terminal subdomain, binds m GTP or cap analogues, positioning the guanine base through π-π stacking with a conserved aromatic residue (Phe/Tyr) and triphosphate stabilization by polar/charged interactions (Lys, Asp, Ser, Asn) that form hydrogen bonds or salt bridges with oxygen of ribose and α-phosphate (Geiss et al., 2009; Liu et al., 2010). This creates a clamp-like fit over the GTP or m G moiety. The sequence and structure of the GTP binding pocket are conserved across mosquito-borne (DENV, ZIKV, JEV) and tick-borne (TBEV, OHFV, LGTV) Orthoflavivirus (Li et al., 2022; Yang et al., 2021; Zhou et al., 2007).

In NS5 MTase, the RNA-binding groove is a shallow, positively charged groove rather than a deep groove (Zhou et al., 2007). This groove lies on the surface of the N-terminal subdomain of MTase, located in close proximity to the SAM and GTP/cap pocket, through which the 5′ viral RNA backbone slides (Zhou et al., 2007). Structural and mutational studies in WNV, DENV (1-4) and related Orthoflaviviruses identify a set of conserved residues that form RNA binding site, including an N-terminal basic cluster (Lys14, Lys17, Lys21-Lys23, Lys29), aromatic and hydrophobic residues near the cap pocket (Phe24/Phe25, Leu17, Leu20), and basic residues along the central groove (Lys41, Lys61, Arg84, Lys105, Lys130, Lys181-Lys183, Lys196 and Lys200 in DENV/WNV-like numbering) (Geiss et al., 2009; Henderson et al., 2011; Liu et al., 2010). The N-terminal helix-turn-helix region of NS5 MTase contains a basic patch (Arg/Lys22, Lys23, Lys28, Lys29, Arg41, Arg42-type positions in DENV/ZIKV), which recognizes and stabilizes the RNA phosphodiester backbone (Henderson et al., 2011).

In the presence of S-adenosylmethionine (SAM), the MTase domain catalyzes the sequential methylation of guanine N-7 and the 2′-OH ribose of genomic RNA cap structure (Zhou et al., 2007). For efficient translation of viral RNA, methylation at the N-7 position is crucial, whereas methylation at 2′-O is pivotal to evading host innate immunity (Chambers et al., 1990; Zhou et al., 2007). The methyltransferase reaction or capping takes place sequentially on an evolutionary conserved orthoflavivirus 5′-RNA terminal (5′AG3′), which serves as the RNA template to produce a mature 5′RNA cap, GpppAG-RNA→m^7^GpppAG-RNA (“cap-0”)→m^7^GpppAm2′-O-G-RNA (“cap-1”) (Cleaves & Dubin, 1979; Zhao, Soh, Lim, et al., 2015). When the 5’ viral RNA arrives, its cap-proximal region binds to this basic groove, bringing the m□G portion of the cap into the GTP/cap pocket and the A1 2’□OH near the SAM pocket. This positioning allows N7 and 2’□O methylation to occur efficiently (Egloff et al., 2002b).

In the absence of MTase activity, plus-strand single-stranded enveloped RNA viruses cannot methylate at the 5′ cap, which affects virus replication by reducing viral genome stability and inefficient translation. Therefore, SAM-dependent viral MTases are potential antiviral targets (Bhutkar et al., 2024, 2025, 2026; Mudgal et al., 2022; Tomar et al., 2011). NS5 MTase methylated RNA suppresses recognition by innate immune sensors of the target cell, which include receptor retinoic acid-inducible gene-I (RIG-I) and interferon-induced protein with tetratricopeptide (IFIT) proteins (Daffis et al., 2010; Zhou et al., 2007). Therefore, NS5 methyltransferase represents an attractive target for antiviral drug design. Structural studies have shown that some viruses, such as DENV NS5 MTase, can form homodimers in a crystal lattice, indicating a potential for protein self-association. (Liu et al., 2010).

Researchers have found various chemical inhibitors which target SAM-dependent viral MTase domains across diverse RNA virus families. Berbamine, venetoclax, and ponatinib have been shown to inhibit the nsP1 MTase of chikungunya virus (CHIKV), resulting in potent suppression of viral replication in cell□based models (Bhutkar et al., 2024). Sinefungin and S□adenosyl□L□homocysteine (SAH), which are SAM analogues, exhibit broad antiviral activity by competitively inhibiting MTases from orthoflaviviruses, alphaviruses, and poxviruses (Chen et al., 2013). In the context of coronaviruses, 3□deazaneplanocin A (DZNep) (Kumar et al., 2022), TDI□015051 (Meyer et al., 2025), HK370 (compound 18l) (Kocek et al., 2024), and NCGC00606183 (Pal et al., 2025) have been reported as antiviral agents that target SARS□CoV□2 MTase activity, mainly through interference with SAM□dependent methylation steps. Ribavirin, a nucleoside analogue, displays multi-viral therapeutics for riboviruses, in part by impairing viral RNA synthesis and cap formation (Beaucourt & Vignuzzi, 2014). Moreover, NSC 111552, NSC 288387 (Samrat et al., 2023), NSC 12155, and NSC 125910 (Brecher et al., 2015) have been characterised as SAM□binding□site inhibitors of orthoflavivirus NS5 MTase, showing suppression of both N^7^ and 2’□O methylation and broad□spectrum antiviral effect in DENV, YFV, and ZIKV. Aurintricarboxylic acid (ATA) has been reported to inhibit Zika and dengue virus NS5 MTase and to exert antiviral activity in cell based experiments (Kaur et al., 2018; Park et al., 2019). Recently, Bhutkar et al.,(2025) identified HC (Herbacetin), a natural polyhydroxylated flavonoid and CAPE (Caffeic acid phenethyl ester), a phenolic antioxidant molecule as potential inhibitors of Dengue NS5 MTase. Additionally, broad-spectrum anti viral activity against Dengue and Chikungunya has been demonstrated for both the molecules HC and CAPE (Bhutkar et al., 2025).

The focus of our present work is the structural and functional characterization of the recombinant NS5 methyltransferase (MTase) domain of KFDV. The recombinant protein has been purified, and crystallised to elucidate the structural basis of its enzyme function. In this investigation the potential antiviral effect of HC and CAPE on the methyltransferase of the tick-borne Orthoflavivirus KFDV has been studied using a combination of biochemical and biophysical techniques.

## 2. Materials and methods

### 2.1 Multiple sequence alignment (MSA) of Orthoflaviviruses NS5 MTase

Kyasanur Forest disease virus (KFDV) full DNA sequence (GenBank: OP037814.1) was retrieved from NCBI nucleotide (Yang et al., 2021). The NS5 MTase domain of KFDV was defined by multiple sequence alignment with other orthoflavivirus NS5 MTase sequences such as ZIKV NS5 MTase (PDB ID-5M5B) (Coutard et al., 2017), DENV-3 NS5 MTase with CAPE and SAH (PDB ID-8KDZ) (Bhutkar et al., 2025), YFV NS5 MTase (PDB ID-3EVA) (Geiss et al., 2009), JEV full length NS5 (PDB ID-4K6M) (Yang et al., 2021), Ntaya virus MTase in complex with GTP and SAH (PDB ID-8CQH) (Krejcova & Boura, 2025) Langat virus MTase (PDB ID-7WNJ) (Li et al., 2022), Tick-borne encephalitis virus MTase (PDB ID-7D6M) (Yang et al., 2021), Omsk hemorrhagic fever virus NS5 MTase (PDB ID-7FJT) (Jia et al., 2022). Multiple sequence alignment was performed and visualized using the multalin tool (Corpet, 1988) and ESPript tool (ENDscript - https://endscript.ibcp.fr), respectively (Robert & Gouet, 2014).

### 2.2 KFDV NS5 MTase cloning, protein expression, and purification

The KFDV NS5 full-length gene was synthesised and received (BIOMATK) in the pET28a+ vector. Using N-terminal His tags, the KFDV NS5 MTase gene (1-813 bp) was cloned into the pET28c+ vector. The amplification of the gene was done using specific forward primer (5′-ATATGCTAGCGGAGGCGCGGAAGGAGAGACACTTGGAGATATTTGG-3′) and reverse primer (5′-ATATCTCGAGCTATACGCAGCGAGTACCCGTGCCCAGATCTACTT-3′) with Phusion DNA Polymerase (New England Biolabs). The PCR amplified product was cloned into the pET 28c+ vector between Xho1 and Nhe1 restriction sites.

The CaCl_2_ heat shock method was used to transform the cloned plasmid into *Escherichia coli* DH5α cells (Yap et al., 2007), and the Plasmid Mini Kit (Qiagen, Hilden, Germany) was used to purify the DNA according to the manufacturer’s protocol. The recombinant plasmid was sequenced to confirm insertion of the NS5 MTase gene into pET 28c+. To express and purify the NS5 MTase protein, the recombinant plasmid with a hexahistidine (6 x His) tag was transformed into *Escherichia coli* BL21 cells. The Luria-Bertani (LB) broth medium containing 50 μg/mL kanamycin was used to culture *Escherichia coli* BL21 bacterial cells. It was incubated at 37 ° C, 180 RPM, until the optical density of 0.6-0.7 at 600 nm (OD 600). The induction of protein expression was done using 0.8 mM of IPTG (isopropyl-β-D-thiogalactopyranoside), and the culture was further incubated for 18 hours at 16°C, shaking at 180 RPM. The culture was harvested after the incubation by centrifugation at 6000 RPM. Using a lysis buffer (50 mM Tris-HCl, 300 mM NaCl, 10% (v/v) glycerol, 20 mM imidazole, pH 8.0), the cells were lysed via French press at 21 kpsi to purify the protein. The lysate was separated by centrifugation for 90 minutes at 12,000 RPM at 4 °C. A nickel-nitrilotriacetic acid (Ni-NTA) column (BioRad, India) was used for purification of 6xHis-tagged KFDV NS5 MTase protein. The column was pre-equilibrated with buffer, and then the supernatant was loaded into the column. The protein was eluted with a linear imidazole gradient in elution buffer [50 mM Tris-HCl, 300 mM NaCl, 10% (v/v) glycerol, 300 mM imidazole, pH 8.0]. The eluted fractions of protein were analysed via 12% SDS-PAGE (sodium dodecyl sulfate-polyacrylamide gel electrophoresis) (Supplementary Fig.1). The eluted protein fractions were used for dialysis, and then, using the amicon centrifugal filters (10000 MWCO; Merck Millipore, Burlington, MA, USA), the protein was concentrated.

**Fig.1.**
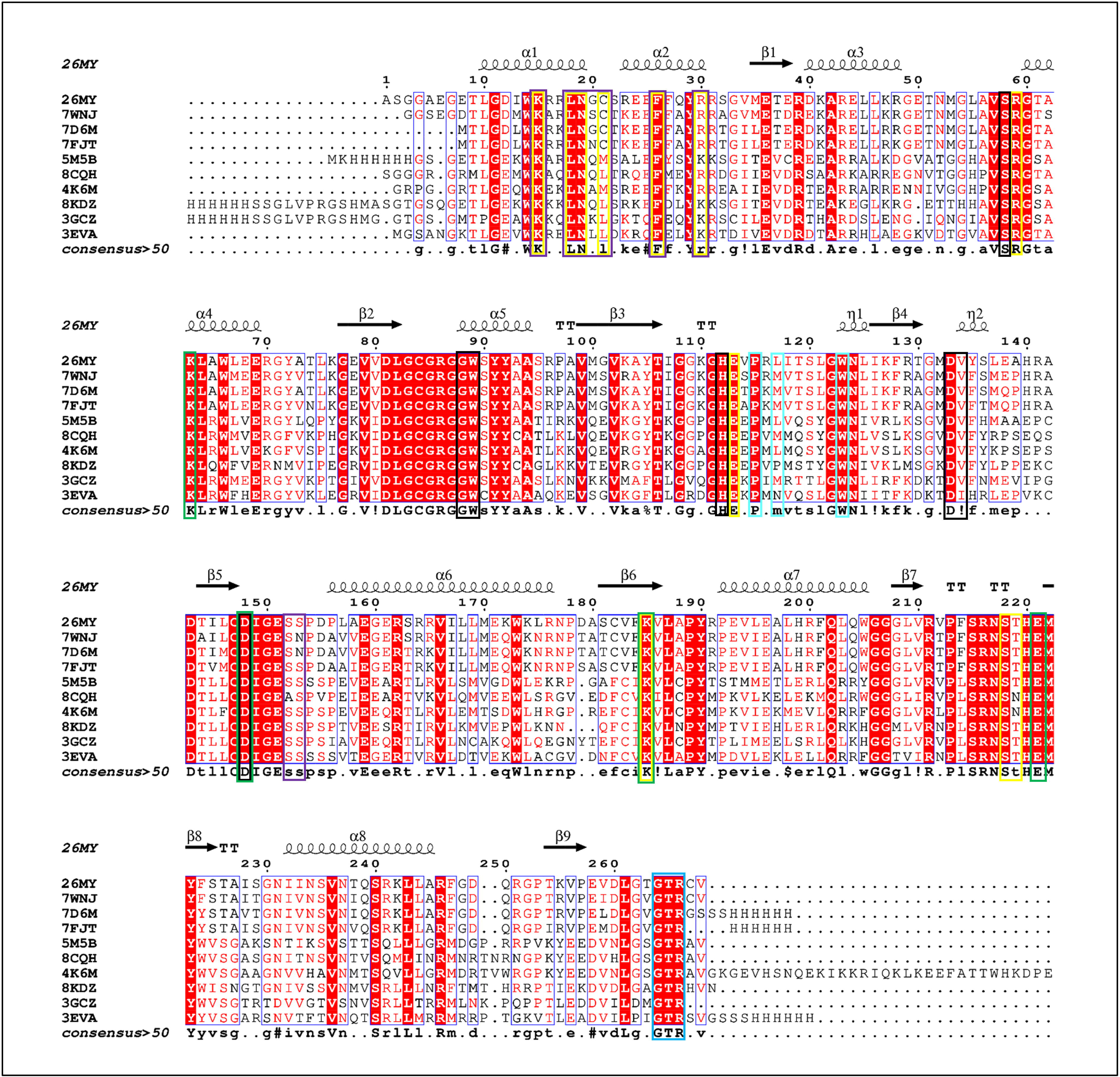
Multiple sequence alignment of Orthoflavivirus NS5 MTase domain-Orthoflavivirus NS5 MTase sequences such as ZIKV NS5 MTase (PDB ID-5M5B), DENV-3 NS5 MTase with CAPE and SAH (PDB ID-8KDZ), YFV NS5 MTase (PDB ID-3EVA), JEV full length NS5 (PDB ID-4K6M), Ntaya virus MTase in complex with GTP and SAH (PDB ID-8CQH) Langat virus MTase (PDB ID-7WNJ), Tick-borne encephalitis virus MTase (PDB ID-7D6M), Omsk hemorrhagic fever virus NS5 MTase (PDB ID-7FJT). MTase catalytic tetrad is shown in green box (K63-D148-K185-E221), C-terminal RdRp interacting GTR motif is shown in blue box (Gly264, Thr265, and Arg266), N-terminal MTase Domain Key-interacting residues with RdRp are shown in cyan box (Pro115, Leu117, Trp123) SAM/SAH binding residue shown in black box (Ser58, Gly88, Trp89, His112, Asp133, Val134, Asp148), GTP Binding Residues shown in purple box (Lys15, Leu18, Asn19, Gly20, Cys21, Phe26, Arg30, Ser152, Ser153) RNA Binding Residues shown in yellow box (Lys15, Leu18, Asn19, Cys21, Phe26, Arg30, Arg60, Glu113/Glu111, Lys185, Ser218, Thr219). Multiple sequence alignment was performed using the multalin tool and ESPript3.2.

### 2.3 KFDV NS5 MTase Crystallization and Data Collection

The crystallization of KFDV NS5-MTase was conducted using the sitting drop vapour diffusion method in 96-well plates at 20°C temperature (Li et al., 2022). After extensive crystallization screening, small rod-shaped crystals were obtained. Diffraction quality crystals were obtained in 6% v/v Tacsimate pH 6.0, 0.1 M MES monohydrate pH 6.0, and 25% w/v PEG 4000. Additionally, crystals were also obtained in 0.2 M Sodium formate, 0.1 M BICINE pH 8.5, 20% w/v PEG 5000 MME (PEG Rx Screen HR2–086, Hampton Research). Larger rod shape crystals were obtained over 5-7 days at 8-10 mg/ml with 0.5 μL of precipitation solution (0.2 M Sodium formate, 0.1 M HEPES, pH 6.5, 20-28% (w/v) PEG 5000).

For data collection, KFDV NS5 MTase crystals in complex with GTP were obtained by soaking the crystals for a few seconds in a cryoprotectant solution containing 30% (v/v) Ethylene glycol (Sigma). The SAH-bound complex was obtained without deliberate co-crystallization or soaking with the co-factor, SAM. The presence of the bound co-factor indicates that it was likely incorporated during heterologous protein expression and retained during subsequent purification and crystallization stages. The crystals were flash-frozen in a nitrogen gas stream at 100 K. The diffraction data of KFDV NS5 MTase was obtained using Rigaku Micromax 007 HF (Tokyo, Japan) at the Macromolecular Crystallography Unit, IIT Roorkee.

### 2.4 Structure determination and refinement

The data reduction was performed using Aimless of the CCP4i2 suite (Potterton et al., 2018). The structure of Orthoflavivirus NS5-MTase (ZIKV NS5 MTase-5M5B) was used as the search model for molecular replacement using the MOLREP program in the CCP4i2 suite. WinCoot and REFMAC5 programs were used for model building and structure refinement, respectively (Emsley et al., 2010; Murshudov et al., 2011). The SAH and GTP molecules were refined with full occupancies. Solvent molecules such as Polyethylene glycol (PEG), ethylene glycol, and water molecules were added where the Fo-Fc map values were above 3σ, and the 2Fo-2Fc map displayed a density at the 1σ contour. The refinement included thirty cycles of rigid-body, followed by restrained refinement, to obtain acceptable *R*_cryst_ and *R*_free_ values. PyMol was used for structural elucidation (The PyMOL Molecular Graphics System, Version 3.0 Schrödinger, LLC). Additionally, composite omit maps were generated to confirm the co-factor and substrate binding in the structure. The data processing and refinement statistics are summarized in Table 1.

**Table 1.** Data processing and refinement statistics for KFDV NS5 MTase.

| Parameter | KFDV NS5 MTase in complex with SAH (26MY) | KFDV NS5 MTase in complex with SAH and GTP (27YN) |
| --- | --- | --- |
| Resolution range | 25.84 - 2.2 (2.279 - 2.2)<sup>a</sup> | 26.42 - 2.6 (2.693 - 2.6)<sup>a</sup> |
| Space group | $P 2_1 2_1 2_1$ | $P 2_1 2_1 2_1$ |
| Unit cell | 48.79 Å 51.67 Å 117.74 Å<br>90.00° 90.00° 90.00° | 49.18 Å 51.86 Å 117.92 Å<br>90.00° 90.00° 90.00° |
| Multiplicity | 5.7 (5.9)<sup>a</sup> | 5.6 (5.9)<sup>a</sup> |
| Completeness (%) | 99.85 (99.87)<sup>a</sup> | 99.76 (99.89)<sup>a</sup> |
| I/$\sigma$(I) | 24.69 (9.89)<sup>a</sup> | 23.73 (9.44)<sup>a</sup> |
| Wilson B-factor (Å<sup>2</sup>) | 15.85 | 19.91 |
| $R_{\text{merge}}$<sup>b</sup> | 0.05918 (0.1739)<sup>a</sup> | 0.06196 (0.1784)<sup>a</sup> |
| $R_{\text{pim}}$ | 0.02676 (0.07689)<sup>a</sup> | 0.02808 (0.07959)<sup>a</sup> |
| CC1/2 | 0.998 (0.981)<sup>a</sup> | 0.998 (0.973)<sup>a</sup> |
| Reflections used in refinement | 15722 (1539)<sup>a</sup> | 9748 (939)<sup>a</sup> |
| Reflections used for R-free | 777 (92)<sup>a</sup> | 479 (48)<sup>a</sup> |
| $R_{\text{work}}$<sup>c</sup> | 0.16 (0.15)<sup>a</sup> | 0.16 (0.16)<sup>a</sup> |
| $R_{\text{free}}$<sup>c</sup> | 0.22 (0.20)<sup>a</sup> | 0.23 (0.28)<sup>a</sup> |
| Number of non-hydrogen atoms | 2243 | 2215 |
| Macromolecules | 2066 | 2066 |
| Ligands | 36 | 69 |
| Solvent | 141 | 80 |
| Protein residues | 259 | 259 |
| RMS (bonds) (Å)<sup>d</sup> | 0.011 | 0.012 |
| RMS (angles) (°)<sup>d</sup> | 2.12 | 2.53 |
| Favored (%) | 98.83 | 98.83 |
| Allowed (%) | 1.17 | 1.17 |
| Outliers (%) | 0 | 0 |
| Rotamer outliers (%) | 1.88 | 2.82 |
| Clash score | 1.9 | 4.23 |
| Average B-factor (Å<sup>2</sup>) | 17.02 | 20.29 |
| Macromolecules (Å<sup>2</sup>) | 16.22 | 19.71 |
| Ligands | 27.64 | 36.99 |
| Solvent (Å<sup>2</sup>) | 25.93 | 20.76 |

**a** Values in parentheses are for the highest resolution shell.
**b** $R_{\text{merge}} = \sum |I - \langle I \rangle| / \sum I$
**c** $R = \sum |F_{\text{obs}}| - |F_{\text{calc}}| / \sum |F_{\text{obs}}|$ . The $R_{\text{free}}$ is the R calculated on the 5% reflections excluded for refinement
**d** RMS is the root mean square

### 2.5 Protein Data Bank accession code

The coordinates and structure factors have been deposited in the Protein Data Bank with accession code 26MY (KFDV NS5 MTase with SAH) and 27YN (KFDV NS5 MTase with SAH and GTP).

### 2.6 HC and CAPE stock solution preparation

Herbacetin (HC) [CAS Number-527-95-7]and caffeic acid phenethyl ester (CAPE) [CAS Number-104594-70-9] were obtained from Cayman Chemical (USA). To prepare stock solutions for experimental use, the compounds were prepared in anhydrous dimethyl sulfoxide (DMSO; Sigma-Aldrich) at final concentrations of 10 mM for HC and 100 mM for CAPE. Followed by storage at -20□°C in aliquots to reduce the detrimental effects of repeated freeze-thaw cycles.

### 2.7 Isothermal Titration Calorimetry and Capillary Electrophoresis-based KFDV NS5 MTase interaction assay

To determine enthalpy (ΔH), dissociation constant (*K_d_*), stoichiometry (n), and entropy (ΔS) of ligands bound with target KFDV NS5 MTase proteins ITC experiment was performed. The binding affinities of the selected ligands (HC and CAPE) were determined using a Microcal-ITC_200_ instrument (Malvern, Northampton, MA) at 25 °C. Initially, the RNA (5’pAGUAA3’ ) and KFDV-MTase protein were dissolved in a DEPC-treated buffer (20 mM HEPES (pH 8.0), MgCl_2_, and 100 mM NaCl) and degassed before ITC experiments. 10-50 μM variable concentrations of KFDV-MTase (in ITC cell) were titrated with 10-100μM range of RNA (in ITC syringe). Likewise, the 10-50 μM of MTase was titrated against a variable range (100-500 μM) of ligands (HC/CAPE). Finally, this saturated MTase-HC/CAPE solution was kept in the ITC cell and titrated against 100 μM RNA (kept in the syringe). The reference cell was filled with water, and the reference power was set to 10 µcal/s. Experiments were performed at a stirring speed of 850 RPM at an initial delay of 100 sec. After the equilibration process, 20 injections were performed; the first injection was 0.4 µl, and the remaining 19 injections (2 µl each) of the syringe samples were titrated into the cell at intervals of 220 sec. The binding isotherm data (n, ΔH, ΔS, and *K_D_* values) were fitted using a one-site binding model and processed using MicroCal Analysis Software (Malvern) and analysed using Origin 7.0 (Dhaka et al., 2025).

The inhibitory activity of HC and CAPE was evaluated against KFDV NS5 MTase, as HC and CAPE have been established as SAM-dependent viral MTase inhibitors (Bhutkar et al., 2024, 2025). The inhibition of KFDV NS5 MTase activity by HC and CAPE was investigated using an *in vitro* biochemical enzyme assay with capillary electrophoresis-based analysis. The reaction mix included 50 mM Tris-HCl (pH 8.0), 2 mM DTT, 10 mM KCl, 2 mM MgCl□, 0.3 mM GTP, 0.3 mM SAM, and 1 μM KFDV NS5 MTase protein. The reactions were incubated for 1 hour at 37 °C, and the activity of the enzyme was measured by the conversion of SAM [*S*-(5′-Adenosyl)-*L*-methionine] to reaction product S adenosylhomocysteine (SAH). Various concentrations of HC and CAPE were tested against the KFDV NS5 MTase protein, and 100 μM caffeine was used as an internal standard.

After incubation, reactions were stopped by the addition of acetonitrile (1:2, v/v). The reaction samples were vortexed for 15 seconds and centrifuged at 18,500 g for 20 minutes to remove precipitated protein, and the supernatants were transferred to autosampler vials to run in an Agilent 7100 CE system (Agilent Technologies, Santa Clara, CA, USA) with diode array detection, using an extended light□path fused□silica capillary (total length 64.5 cm, effective length 56 cm, internal diameter 50 μM). It is operated at 30 kV, 20°C capillary temperature, and 8°C tray temperature. Hydrodynamically, samples were injected at 50 mbar for 10 seconds. As previously mentioned in Mudgal et al. 2020, for every experiment, the capillary was pre-conditioned using a 5-minute CE-grade water rinse, a 10-minute NaOH rinse, a 5-minute water rinse, and a 10-minute running buffer rinse. Rinsed between injections and flushed with water, followed by buffer wash. SAH peak areas were normalised to caffeine, as caffeine was run as an internal control. Data analysis from three experiments was done with GraphPad Prism 8 using nonlinear regression with a “one-site inhibition model” to determine the inhibition parameters of HC and CAPE (Bhutkar et al., 2024, 2025; Mudgal et al., 2020).

### 2.8. RNA binding inhibition using fluorescence polarization (FP) assay

The fluorescence polarisation (FP) assay was used to analyse KFDV NS5 MTase protein and FAM-labelled RNA binding inhibition by small inhibitory molecules HC and CAPE. Previously established FP assay used for studying SARS-CoV2 NP protein and RNA binding was used with small modification (Dhaka et al., 2025). In brief, the 5-mer fluorescent FAM-labelled RNA probe [5’pAGUAA-FAM 3’] was chemically synthesised and purchased from Biotech Desk Pvt Ltd (India). To maintain RNase-free environment the DEPC-treated water was used to prepare RNA stock, and buffers solutions. All reactions were prepared in 50mM Tris-HCl, 10mM KCl, 2mM MgCl_2_, 2mM DTT (pH=8.0). KFDV NS5 MTase was used at a final concentration of 2.5 µM in each well, and FAM-labelled RNA was used as the fluorescent ligand, as we are targeting RNA binding site of the protein. The compounds herbacetin (HC) and caffeic acid phenethyl ester (CAPE) were used at various concentrations to assess their inhibitory effect on KFDV NS5 MTase protein-RNA complex formation. These assays were set up in a 96-well black, flat-bottom polystyrene NBS microplate (Corning, Cat. No. 3686; nonsterile, without lid), with a total reaction volume of 100 µl per well, in triplicate at room temperature. The Fluorescence polarisation was measured by using a green filter set, with excitation at 485/20 nm and emission at 528/20 nm. This experiment was done by Synergy H1 multimode plate reader (Agilent BioTek). GraphPad Prism 8 software was used for data analysis. Binding and inhibition curves were fitted by nonlinear regression using one site-specific binding.

### 2.9 Tryptophan fluorescence spectroscopy assay

The tryptophan fluorescence spectroscopy (TFS) is a label-free, valuable technique used to investigate protein-ligand interactions by measuring the emission of tryptophan residues of the protein. TFS has been previously used Bhutkar et al., 2025 to study binding of ligands to orthoflavivirus NS5 MTase (Bhutkar et al., 2025). The protein samples were diluted in 1x PBS (Phosphate-Buffered Saline) buffer, and each well contained 2.0 µM KFDV NS5 MTase protein. In the same buffer conditions, small inhibitory compounds, HC and CAPE, were used to check binding with KFDV MTase protein. GTP and 5’pAGUAA3’ RNA were used as ligands to target RNA-binding site of protein. Tryptophan fluorescence spectroscopy 96-well black, flat-bottom polystyrene NBS microplates (Corning, Cat. No. 3686; non-sterile, non-lid) were utilised, with 100 µl sample per well. The experiment was done at room temperature (∼25 °C), with an excitation at 280 nm, and emissions from 300 to 600 nm. The data was analysed by GraphPad Prism 8 software. Binding isotherms and inhibitor titration curves were fitted through nonlinear regression with “One Site–Specific Binding” model. The Synergy H1 multimode plate reader (Agilent BioTek) was utilized to measure the change in intrinsic tryptophan fluorescence of KFDV NS5 MTase protein, to observe conformational changes after ligand binding.

## 3. Results

### 3.1 Multiple Sequence Alignment Reveals Conservation of the Methyltransferase Active Site and Key Functional Residues Across Orthoflaviviruses

The multiple sequence alignment of Orthoflavivirus NS5 MTase domains, based on crystal structure PDB IDs 7WNJ, 7D6M, 7FJT, 5M5B, 8CQH, 4K6M, 8KDZ, 3GCZ and 3EVA, revealed highly conserved SAM binding, RNA binding, and GTP binding site amino acid residues, including KFDV NS5 MTase PDB ID 26MY. The conserved K–D–K–E catalytic motif was highly conserved across all MTase amino acid sequences. It confirms a common catalytic mechanism for N7 and 2’O methylation across all Orthoflaviviruses. The residues that are important for the GTP or cap-binding pocket in cap-bound structures were also well conserved. The aromatic residues (Phe-24) that stack with the guanine base of RNA exhibited strong conservation across the alignment. The RNA-binding site residues are also conserved across the NS5 MTases (Fig. 1).

### 3.2 KFDV NS5 MTase protein purification

KFDV NS5 MTase protein was expressed in *Escherichia coli* BL21 cells and purified using Ni-NTA affinity chromatography, it was followed by size-exclusion chromatography (SEC). SDS-PAGE analysis of purified protein showed a single dominant band at approximately ∼30 kDa, indicating high purity of the protein (Supplementary Fig. 2).

### 3.3 Structural analysis of KFDV NS5 MTase protein

#### 3.3.1 Binary complex of KFDV NS5 MTase and S-adenosyl-L-homocysteine (SAH)

KFDV NS5 MTase domain present at N-terminus of NS5 displays a conserved SAM-dependent MTase fold characterized by the presence of a classical Rossman fold, which is crucial for the binding of SAM co-factor. With a total of 265 residues, the MTase domain of KFDV consists of nine α helices and nine β-strands forming a central β-sheet surrounded by α-helices on both sides. The central β-sheet is constituted by strands β2-β8. The Rossmann fold creates a deep cleft between the β-sheet and the flanking helices, which serves as the SAM-binding pocket. This pocket is lined by critical interacting residues that stabilize the product, i.e. SAH molecule via hydrogen bonding and Van der Waals’ interactions. The OXT (C-terminal oxygen atom) of SAH interacts with Ser58 and Gly88 via conventional hydrogen bonds at 2.7 and 3.0 Å, respectively, while Asp148 and N atom (at 2.9 Å), His112 and O3’(at 2.9 Å), Asp133 and N6 (at 3.1 Å), Val134 and N1 (at 3.0 Å) also show major stabilizing H-bonds. The adenine ring of the SAH is stabilized by Ile107 and Ile149 via pi-alkyl and pi-sigma bonds, respectively. A series of water molecules, namely HOH 25, 27, 45, and 53, stabilize the SAH molecule via an H-bonding network. Apart from these critical residues and water molecules, the SAH also makes molecular contacts via van der Waals forces with Gly60, Gly63, Gly83, Cys84, Arg86, Gly87, Trp89, Glu113, Met132, and Tyr135.

The active site is positioned at the interface between the central β-sheet and surrounding α-helices, creating a binding pocket that accommodates both the substrate SAM and the RNA substrate. The catalytic core includes conserved K-D-K-E tetrad (Lys63, Asp148, Lys185, Glu221) located in motifs I, II, and III, which are critical for SAM binding and catalytic activity, with key catalytic residues including Lys63 (motif I), Asp148 (motif II), Lys185 (motif III), and Glu221 (motif IV) forming a conserved network essential for methyl transfer activity. This K-D-K-E tetrad creates a positively charged environment that stabilizes the transition state during methyl transfer to the RNA cap substrate. Motif I (GXGXXG) forms the SAM cofactor binding loop, while motif II contains the catalytic Lys185 residue responsible for methyl transfer (Chouhan et al., 2019).

#### 3.3.2 Ternary complex of KFDV NS5 MTase and Guanosine-5’-triphosphate (GTP)

The RNA-binding site is situated on the opposite side of the SAM-binding pocket and is characterized by a groove rich in basic residues that interact with the RNA substrate’s phosphate backbone. This region includes loops connecting β-strands β7-β8 and β8-β9, which form a flexible RNA-binding interface that accommodates the 5’-terminal RNA cap structure (GpppA-RNA). The GTP molecule, when soaked with KFDV MTase apo crystals in the mother liquor, yielded a GTP-bound complex since the RNA cap is derived from GTP. This complex is critical for gaining fundamental insights into the interaction between protein and RNA cap. The O6 of GTP forms an H-bond with Arg23 (at 2.9 Å), N2 forms an H-bond with Leu18 (at 2.8 Å) and Cys21 (at 3.0 Å), O2’ forms an H-bond with Lys15 (at 2.7 Å) and Asn19 (at 2.6 Å), O2G and O3G form H-bonds with Ser152 (at 2.6 and 2.8 Å, respectively). O2A and O2B from the phosphate moiety form critical salt bridges with Arg30 (at 2.2 and 3.3 Å, respectively) and Arg216 (at 3.0 Å). The guanine ring is stabilized by π-π stacking with Phe26. Apart from these crucial residues, the GTP molecule is also stabilized by Ser218, Ser153, Pro154, Leu187, Glu150, Gly151, and Thr219 via van der Waals forces. This complex also contains a bound SAH molecule, which is stabilized by similar interactions as described above; however, the methionyl tail of SAH, particularly the carboxylate moiety, interacts differently in the GTP-SAM ternary complex. Asp148 makes an H-bond with N of SAH at 2.9 Å, while the same residue forms an H-bond with carboxylic OXT at 3.0 Å. The Cys84 residue, which forms a van der Waals interaction in the apo structure, forms an H-bond with N at 3.4 Å in the GTP-bound complex. Similarly, Trp89 forms a van der Waals interaction in the apo, while it forms an H-bond with carboxylic O at 3.0 Å. Ser58 continues to form an H-bond with OXT at 2.7 Å in the apo while forming the same interaction with N at 3.3 Å in the GTP complex. Gly88 participates in H-bonding via the backbone amino group to the OXT of SAH at 3.0 Å in the apo, while it becomes distant and stabilises the SAH molecule via van der Waals’ interactions in the GTP complex. (Fig.2B)

**Fig.2.**
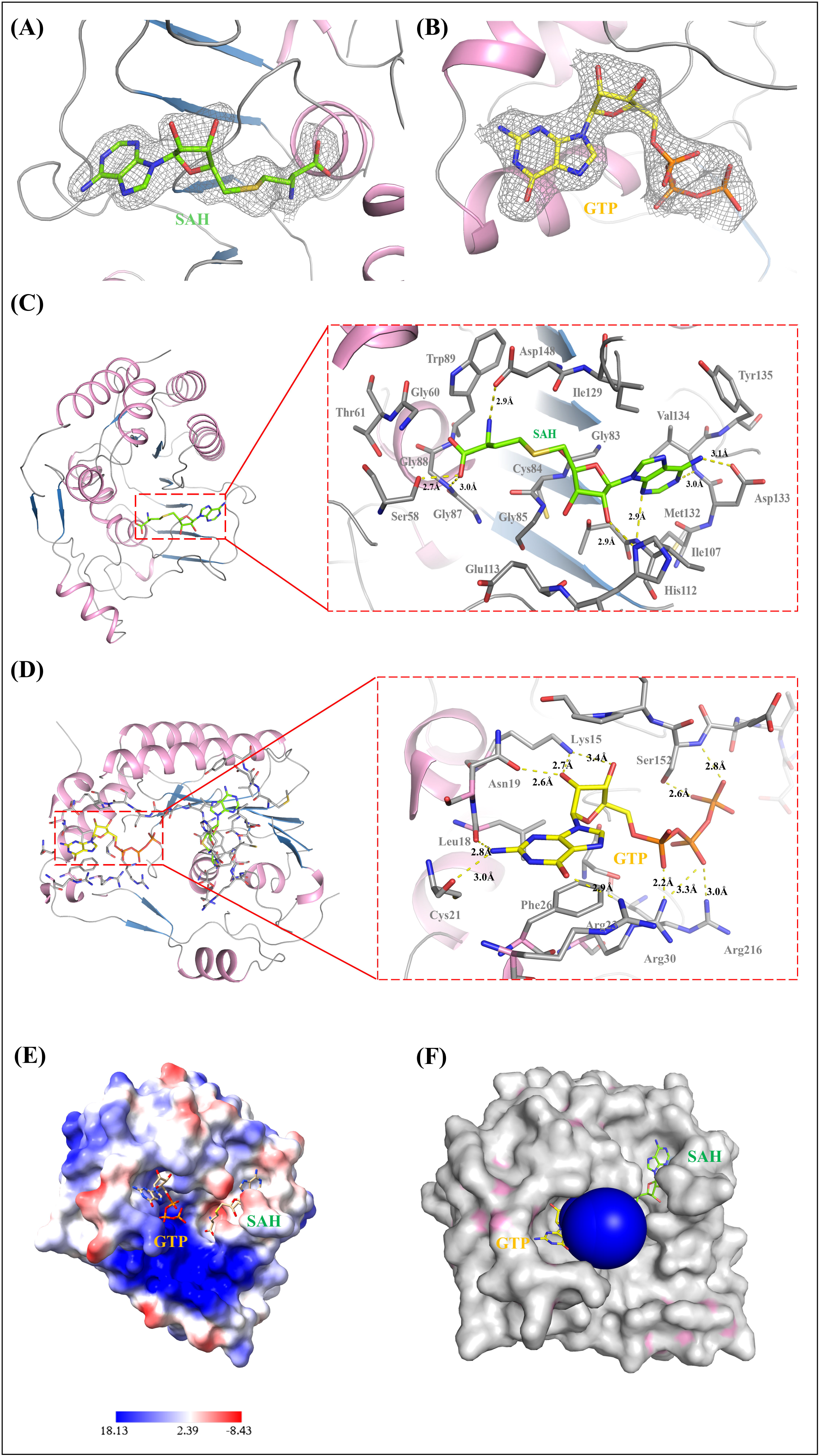
Cartoon representation of KFDV NS5 MTase with bound (A) GTP and (B) SAH showing an electron density Fo-Fc composite omit map (contoured at 3σ, at a carve level of 1.6 Å). (C) The KFDV NS5 MTase-SAH complex structure depicts critical residues that stabilize the cofactor in the binding pocket. The OXT (C-terminal oxygen atom) of SAH interacts with Ser58 and Gly88 via conventional hydrogen bonds at 2.7 and 3.0 Å, respectively, while Asp148 and N atom (at 2.9 Å), His112 and O3’(at 2.9 Å), Asp133 and N6 (at 3.1 Å), Val134 and N1 (at 3.0 Å) also show major stabilizing H-bonds. The adenine ring of the SAH is stabilized by Ile107 and Ile149 via pi-alkyl and pi-sigma bonds, respectively. (D) The KFDV NS5 MTase-GTP complex structure depicts critical residues that stabilize the RNA cap in the active site. The GTP molecule’s O6 forms an H-bond with Arg23 (at 2.9 Å), N2 forms an H-bond with Leu18 (at 2.8 Å) and Cys21 (at 3.0 Å), O2’ forms an H-bond with Lys15 (at 2.7 Å) and Asn19 (at 2.6 Å), O2G and O3G form H-bonds with Ser152 (at 2.6 and 2.8 Å, respectively). O2A and O2B form critical salt bridges with Arg30 (at 2.2 and 3.3 Å, respectively) and Arg216 (at 3.0 Å). The guanine ring is stabilized by pi-pi stacking with Phe26. (E) Electrostatic surface representation of the protein bound to GTP and SAH. The surface is colored by electrostatic potential, with blue indicating positively charged regions and red indicating negatively charged regions. The ligands are displayed as sticks. (F) The tunnel analysis of the protein revealed a major tunnel leading to the RNA-binding site on its surface. The protein has been depicted in surface, and the GTP and SAH are depicted as sticks.

The analysis of the electrostatic surface of the KFDV NS5 MTase protein revealed non-uniform charge distribution around the ligand-binding cleft, with a predominantly positive electrostatic environment surrounding the bound GTP and a neutral to partially negatively charged pocket accommodating SAH. This arrangement is consistent with structural studies of other Orthoflavivirus methyltransferases. In our structure, the bound ligands occupy adjacent but distinct pockets on the protein surface, suggesting that the local electrostatic topology is compatible with the recognition of the substrate and the co-factor. This charge distribution helps bind the negatively charged phosphate moieties and orient the ligands for catalysis. The tunnel analysis performed with CAVER 3.0.3 indicated a single major tunnel originating from the GTP-binding site, with a throughput of 0.95, a bottleneck radius of 3.53 Å, a length of 4.99 Å, and a curvature of 1.37 Å (Chovancova et al., 2012; Kayastha et al., 2026). These parameters indicate that the GTP-binding site is rather exposed and on the protein’s surface, making it readily accessible to the RNA.

#### 3.3.3 Comparison of KFDV MTase with homologous Orthoflavivirus MTases

To assess the conservation of ligand recognition within Orthoflavivirus methyltransferases, the SAM and GTP-bound structures of KFDV NS5 MTase were compared with several homologous viral MTases. Despite sequence divergence among these viruses, the overall architecture of the ligand-binding pockets and the residues involved in ligand stabilization are highly conserved, consistent with the low structural RMSD values observed (0.400 - 0.666 Å) (Fig.3).

**Fig.3.**
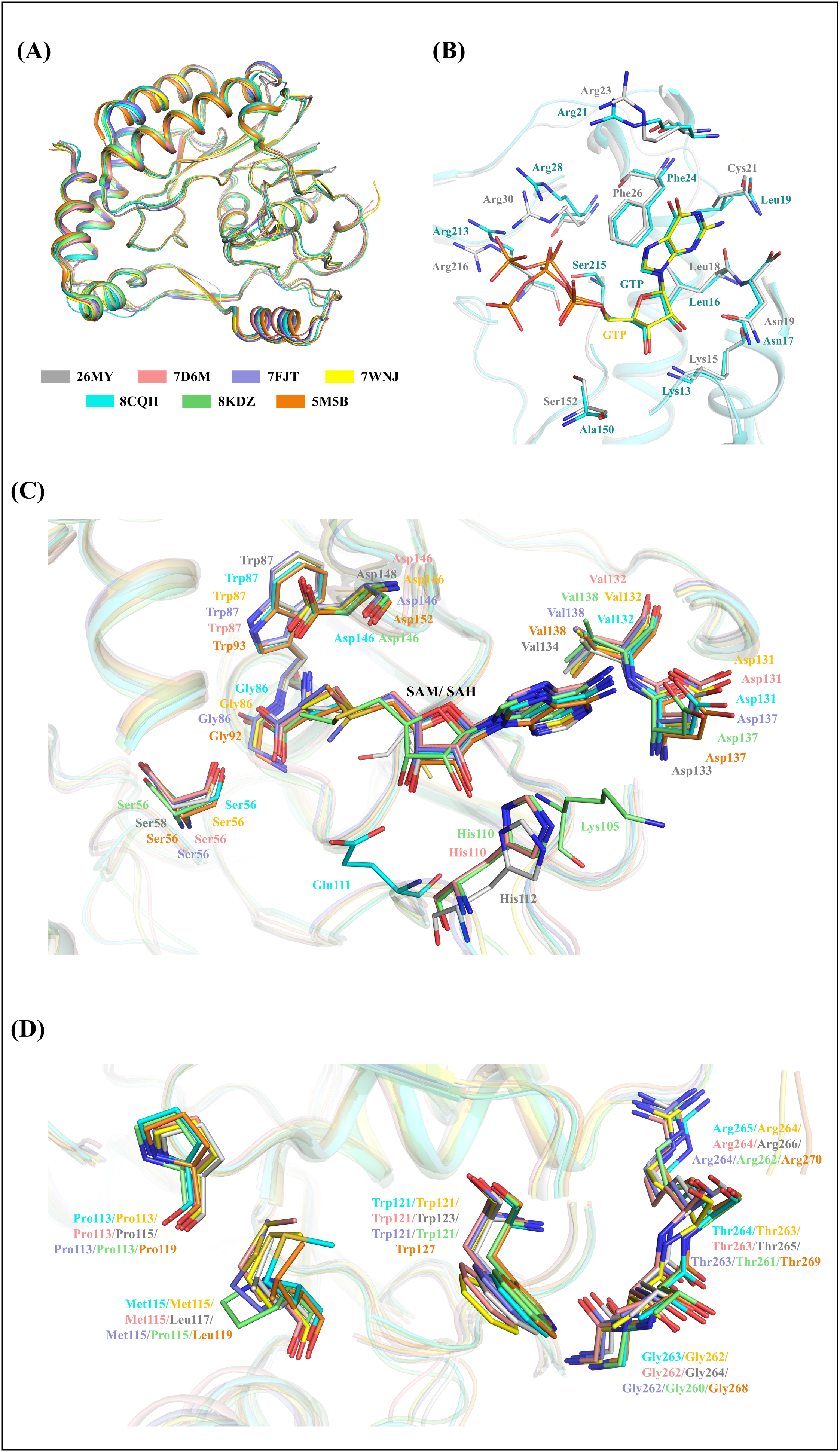
(A) The superposition of KFDV NS5 MTase (26MY) with the homologous proteins Langat virus NS5 MTase (7WNJ, RMSD 0.400 Å), Tick-borne encephalitis virus NS5 MTase (7D6M, RMSD 0.473 Å), Omsk hemorrhagic fever virus MTase (7FJT, RMSD 0.481 Å) , Dengue virus type 3 MTase (8KDZ, RMSD 0.666 Å). Zika virus MTase (5M5B, RMSD 0.580 Å), Ntaya virus MTase (8CQH, RMSD 0.571 Å) suggest some minor differences in the loop and terminal regions (B) the RNA-cap binding region, when compared between 26MY and 8CQH displays similar binding interaction for GTP with some differences observed in the orientation of phosphate moiety of GTP. (C) SAM binding site also indicates a conserved set of interacting residues with minor differences among homologues, that stabilize the SAM co-factor in its binding site. (D) The residues that are critical for the interaction between MTase and RdRp have been compared among the homologues and have been observed to be conserved. However, 26MY and 5M5B have a Leu residue and 8KDZ has a Pro residue instead of the consensus Met residue present in the most homologues.

In KFDV MTase, a network of hydrogen bonds stabilises the SAH, which involves His112, Asp133, Val134, Asp148, Gly88, and Ser58, as well as water-mediated interactions and hydrophobic contacts from Leu107, Ile149, and Met132. Comparison with homologous structures revealed remarkable conservation of these interactions. The closest structural homolog, Langat virus MTase (7WNJ, RMSD 0.400 Å), displays nearly identical SAH recognition, stabilized by hydrogen bonds involving Asp131, Val132, Asp146, Trp87, Ser56, and Gly86, while hydrophobic interactions are mediated by Ile105 and Ile147. These residues correspond directly to Asp133, Val134, Asp148, Trp89, Ser58, Gly88, Leu107, and Ile149 in KFDV, indicating a highly conserved SAH-binding environment. Similarly, Tick-borne encephalitis virus MTase (7D6M, RMSD 0.473 Å) exhibits a nearly identical interaction pattern. Hydrogen bonds are formed with His110, Asp131, Val132, Asp146, Trp87, and Ser56, while Ile105 and Ile147 contribute hydrophobic interactions with the adenine ring. Most residues involved in ligand stabilization are conserved in KFDV, demonstrating strong preservation of the catalytic pocket among tick-borne flaviviruses. The Omsk hemorrhagic fever virus MTase (7FJT, RMSD 0.481 Å) contains a bound SAM molecule. The key residues stabilising the interaction are Ser56, Thr59, Trp87, Thr104, Ile105, His110, Glu111, Asp131, Phe133, and Ile147 correspond directly to Ser58, Thr61, Trp89, Thr106, Leu107, His112, Glu113, Asp133, Tyr135, and Ile149 in KFDV. The observed conservation highlights the recognition of SAM in tick-borne flaviviruses to be governed by similar residues. A similar binding mode is observed in Dengue virus type 3 MTase (8KDZ, RMSD 0.666 Å). Although the overall interaction network is highly conserved, one notable variation occurs at position 105, where Lys105 replaces the hydrophobic Ile or Leu observed in KFDV and most other homologs. Consequently, Lys105 directly participates in hydrogen-bonding and hydrophobic interactions with SAH. Despite this substitution, the overall architecture of the binding pocket remains unchanged, preserving ligand stabilization through conserved residues including Ser56, Ser59, Trp87, Thr104, His110, Glu111, Asp131, Phe133, and Ile147.

The Zika virus MTase (5M5B, RMSD 0.580 Å) contains bound SAM and exhibits strong conservation of the SAM-binding site despite differences in residue numbering. Key interacting residues include Ser62, Ser65, Trp93, Thr110, Lys111, His116, Glu117, Asp137, Phe139, and Ile153, which correspond structurally to Ser58, Thr61, Trp89, Thr106, Leu107, His112, Glu113, Asp133, Tyr135, and Ile149 in KFDV. Similar to the dengue virus, a lysine residue occupies the position corresponding to Leu107 or Ile105 in tick-borne flaviviruses, indicating an alternative but functionally equivalent mode of adenine-ring stabilization. The Ntaya virus MTase (8CQH, RMSD 0.571 Å) exhibits an interaction network almost identical to that of KFDV MTase. SAH is stabilized through hydrogen bonds with Ser56, Gly86, Trp87, Glu111, Asp131, Val132, Asp146, and, while Lys105 and Ile147 provide hydrophobic interactions with the adenine moiety. The extensive conservation of these interactions further supports a common mechanism of SAH recognition across Orthoflavivirus MTases.

Overall, comparison of all SAH or SAM-bound homologs reveals a highly conserved catalytic pocket centred around the residues corresponding to Ser58, Thr61, Trp89, Thr106, His112, Glu113, Asp133, Tyr or Phe135, and Ile149 in KFDV MTase. These residues form the integral SAM-recognition network and may be essential for methyltransferase function. KFDV MTase possesses a bound GTP molecule, stabilized by hydrogen bonds with Lys15, Leu18, Asn19, Cys21, Arg23, and Ser152, as well as electrostatic interactions involving Arg30 and Arg216. The guanine base is further stabilized by π–π stacking with Phe26, while Pro154 and Thr219 contribute additional carbon-hydrogen bonding interactions. Among the homologous structures examined, only Ntaya virus MTase (8CQH) contains both SAH and GTP. Strikingly, the GTP-binding mode is highly conserved between Ntaya virus and KFDV MTase. In the Ntaya virus, GTP is stabilized by hydrogen bonds involving Lys13, Leu16, Asn17, Leu19, Arg28, Arg213, and Ser215, residues that correspond closely to Lys15, Leu18, Asn19, Arg30, Ser152, and Arg216 in KFDV. In both structures, the phosphate groups are stabilized by positively charged arginine residues and extensive water-mediated interactions. The presence of a Mg² ion coordinated to the triphosphate moiety through a network of water molecules is a notable feature of the Ntaya virus structure. Although a magnesium ion is not observed in KFDV NS5 MTase protein, the surrounding residues and overall binding geometry are highly similar, suggesting that both proteins employ a conserved mechanism for nucleotide stabilization. Furthermore, the guanine base participates in π–π stacking with Phe24 in Ntaya virus, equivalent to the Phe26 mediated stacking interaction observed in KFDV MTase. The differences in the orientation of the phosphate moiety of GTP were observed in both structures, where the oxygen moieties of the triphosphate groups are stabilized via salt bridges by Arg30, Arg216, and via H-bonds by Ser152 in the case of KFDV MTase, whereas in Ntaya MTase, Arg28, Arg213, and Ser215 stabilize the same groups via H-bonds. Arg30 of KFDV superimposes with Arg28 of Ntaya MTase, while the other two interacting residues are distantly located.

Despite originating from diverse tick-borne and mosquito-borne flaviviruses, all homologous MTases examined display highly conserved SAH/SAM- and GTP-binding pockets. The low RMSD values and preservation of key catalytic residues indicate that the ligand-recognition mechanism is evolutionarily conserved across flaviviruses, supporting a common mechanism of methyl donor and nucleotide recognition.

Based on structural superimposition, KFDV NS5 MTase may also interact with the finger domain of RdRp via critical hydrophobic contacts involving Pro115, Leu117, and Trp123. These residues correspond to Pro113, Leu115, and Trp121 in orthoflavivirus MTases such as LGTV (PDB ID: 7WNJ) and JEV (PDB ID: 4K6M), known to interact with the RdRp finger domain. 26MY has Leu117, 5M5B has Leu119, and 8KDZ has Pro115 at this position; however, MSA indicates a Methionine residue as the consensus at this position. Another set of residues at the N-terminus, Gly264, Thr265, and Arg266, is typically conserved among related MTases (Krejčová et al., 2024).

### 3.4 KFDV NS5 MTase inhibition and binding to HC and CAPE

HC and CAPE have been established as SAM-dependent NS5 MTase inhibitors, and co-crystal structure of both molecules in complex with Dengue NS5 MTase indicates binding of these compounds to GTP and RNA binding pockets. The thermodynamic binding titration experiments were performed using a MicroCal ITC200 microcalorimeter (Malvern, Northampton, MA) at 25 °C, with variable concentrations of selected ligands HC and CAPE titrated against 30 μM KFDV NS5 MTase protein (in cell) and yielded *K_d_*values of 35 μM and 105 μM for HC and CAPE, respectively. The data were analyzed using Origin 7.0, and the thermodynamic parameters are shown in detail (Fig.4 and Table 2). Enzymatic activity of purified KFDV NS5 MTase protein was evaluated using capillary electrophoresis (CE) in the presence of HC and CAPE. CE assay evaluated the conversion of SAM to SAH, and 1mM caffeine was used as an internal control, and normalised peak areas were used to quantify enzymatic activity. In the presence of HC and CAPE, a significant decrease in SAH formation was monitored, which confirmed the inhibition of KFDV NS5 MTase.

**Fig.4.**
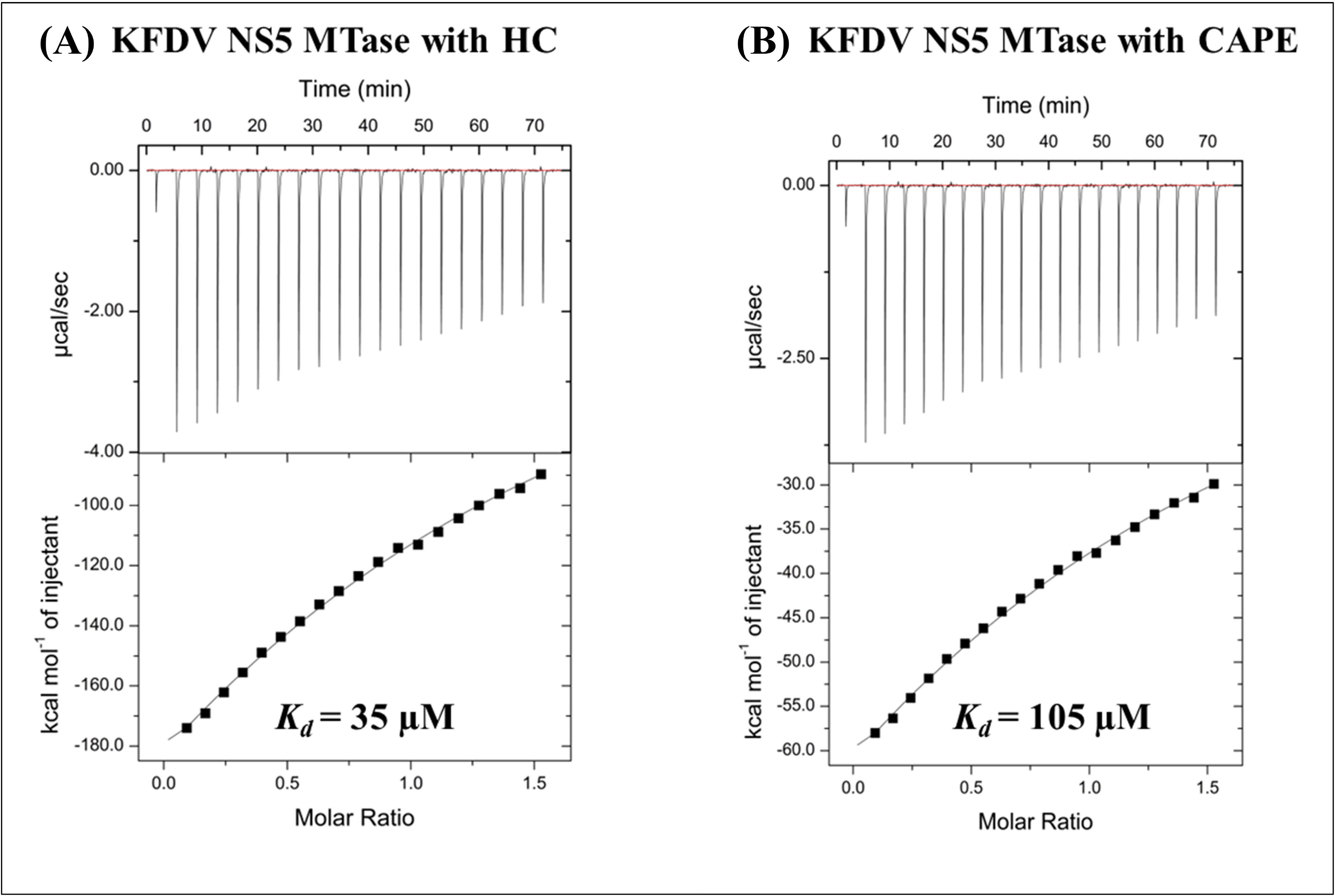
Isothermal titration calorimetry (ITC) analysis of interaction of KFDV NS5 MTase with HC and CAPE. Representative raw thermograms showing the heat response generated by sequential injections of (A) HC and (B) CAPE into the protein. The calculated dissociation constants (K_d_) were **35** μ**M** for compound HC and **105** μ**M** for compound CAPE, indicating their binding affinities for KFDV NS5 MTase protein.

**Table 2.** The thermodynamic analysis of selected ligands (RNA/HC/CAPE) with KFDV-MTase protein, as obtained from ITC.

| ITC-cell | Syringe | n | $K_d$<br>(µM) | $K_A$<br>(M<sup>-1</sup>) | $\Delta H$<br>(cal/mol) | $\Delta S$<br>(cal/mol/degree) |
| --- | --- | --- | --- | --- | --- | --- |
| MTase | RNA | 1 | 15 | $(6.4 \pm 1.8)10^4$ | $(-1.8)10^6 \pm (2.3)10^6$ | $(-6.1)10^3$ |
| MTase | HC | 1 | 35 | $(2.8 \pm 0.7)10^4$ | $(-8.0)10^5 \pm (1.4)10^4$ | $(-2.6)10^3$ |
| MTase-HC<br>saturated | RNA | 1 | 256 | $(3.9 \pm 0.8)10^3$ | $(-3.8)10^6 \pm (6.1)10^4$ | $(-1.3)10^4$ |
| MTase | CAPE | 1 | 105 | $(9.4 \pm 0.2)10^3$ | $(-2.6)10^5 \pm (0.4)10^4$ | $(-0.8)10^3$ |
| MTase-CAPE<br>saturated | RNA | 1 | 344 | $(2.9 \pm 0.3)10^3$ | $(-8.9)10^6 \pm (1.0)10^6$ | $(-3.0)10^4$ |

### 3.5 KFDV NS5 MTase HC and CAPE inhibiting RNA binding using ITC

The thermodynamic binding titration experiments were performed to determine RNA binding with KFDV NS5 MTase protein, using MicroCal ITC_200_ microcalorimeter (Malvern, Northampton, MA) at 25□°C. Binding saturation was observed upon titrating 30 μM of KFDV NS5 MTase (in ITC cell) with 100 μM of RNA (in syringe), and a K_D_ (15 μM) was obtained. After this, a saturated solution of this protein-ligand (in cell) was titrated with the finalised RNA (100 μM) and obtained the K_d_ = 256 μM, and 344 μM, for MTase-HC and MTase-CAPE, respectively (Table 1). The ITC-analysed dissociation constants (K_d_) are shown in detail (Fig.5 and Table 2). The results suggested that the selected inhibitors weakened KFDV NS5 MTase-RNA binding, as evidenced by decreased interactions by ∼17-fold in the presence of HC and ∼22-fold for CAPE.

**Fig.5.**
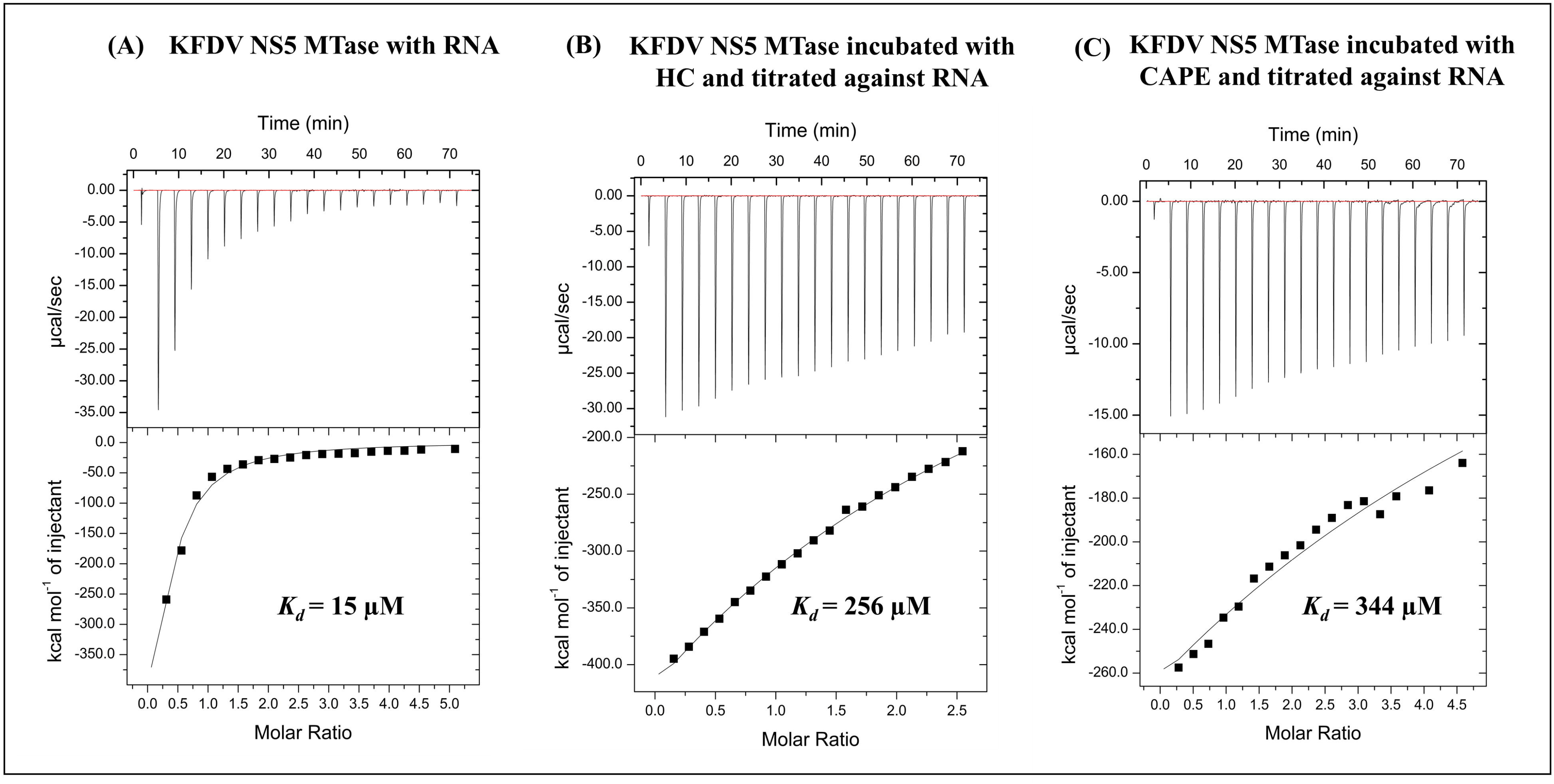
Effect of HC and CAPE on the binding of RNA to KFDV NS5 MTase protein measured by ITC. The ITC experiment shows (A) Direct binding of RNA with KFDV NS5 MTase protein in absence of HC and CAPE.[K_d_=15μM] (B) Binding of RNA to KFDV NS5 MTase protein pre-incubated with HC [K_d_=256μM]. (C) Binding of RNA to KFDV NS5 MTase protein pre-incubated with CAPE [K_d_=344μM]. The increased K_d_, values indicate reduced apparent affinity of RNA after pre-binding of compounds HC or CAPE.

### 3.6 Fluorescence Polarization (FP) based assay of HC and CAPE interaction with KFDV NS5 MTase

To analyse interaction between KFDV MTase protein and FAM-labelled phosphate-modified RNA, fluorescence polarisation (FP) assays were conducted using the 5-mer fluorescent FAM-labelled RNA probe. The FP assay helped to determine binding affinity of KFDV NS5 MTase to the FAM-labelled phosphate-modified RNA probe. To find the optimal RNA concentration, a concentration-response experiment was carried out by incubating FAM-labelled RNA (0.1-1.0 μM) with a fixed concentration of KFDV MTase (10 μM). The binding profiles indicated that 100 nM RNA is suitable for subsequent assays (Fig.6A).

**Fig.6.**
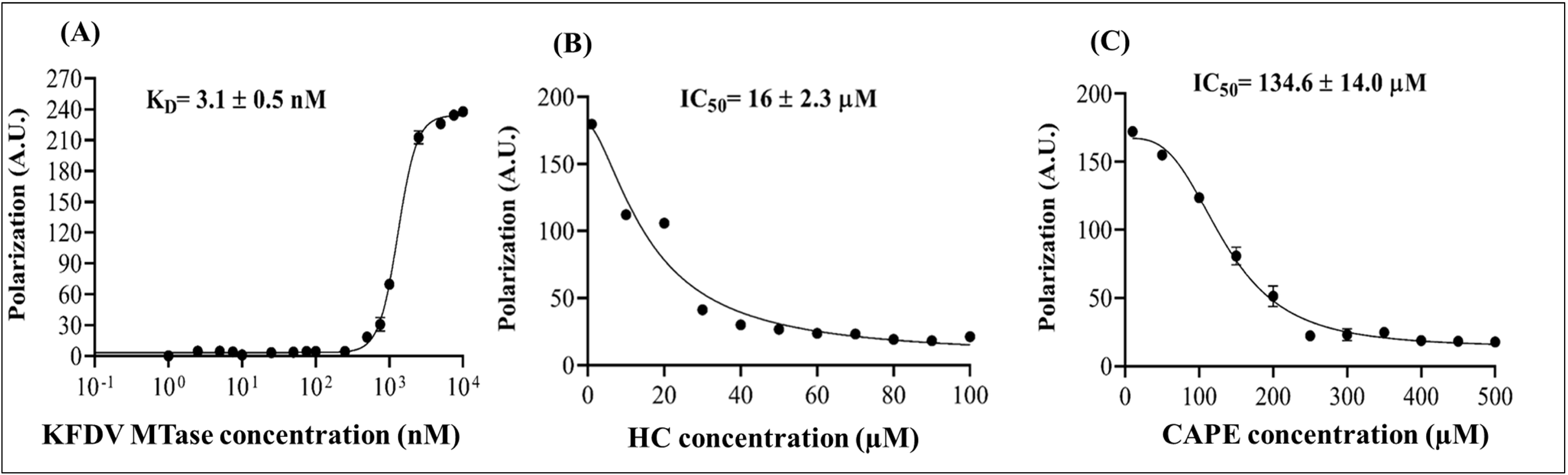
Fluorescence Polarization (FP) based assay of HC and CAPE interaction with KFDV NS5 MTase. (**A**) A concentration-dependent increase in polarization shows interaction of KFDV NS5 MTase with 5’ 6-FAM-labelled RNA. The FP signal response illustrating inhibitory effects of increasing concentrations of (**B**) HC compound, (**C**) CAPE compound for MTase-RNA binding. The standard deviation (SD) was calculated using triplicate values and represent as error bars.

The purified KFDV NS5 MTase protein, ranging from 1 nM to 10000 nM, was prepared in 10 mM KCl, 50 mM Tris (pH 8.0), 2 mM MgCl_2_, and 2 mM DTT buffer. Each concentration of protein was mixed with 100nM FAM-RNA in a 96-well black flat-bottom polystyrene NBS Microplates (nonsterile, without lid, from Corning). Synergy H1 multimode plate reader (Agilent BioTek) was utilized to collect FP assay data with excitation at 480 nm and emission at 535 nm. Following the affinity assessment of RNA binding to the KFDV NS5 MTase protein, fluorescence polarisation (FP) analysis was performed with HC and CAPE, known inhibitors of DENV NS5 MTase. Serial dilutions of HC (from 1 μM to 100 μM) and CAPE (from 10 μM to 500 μM) were diluted in buffer solution containing 10 mM KCl, 50 mM Tris (pH 8.0), 2 mM MgCl_2_, and 2 mM DTT. For inhibition testing, a final concentration of 2.5 μM KFDV NS5 MTase protein and 100 nM RNA were used, with compounds at various dilutions, and incubated for 60 minutes. The FP measurements showed a significant decrease in RNA binding affinity to the KFDV NS5 MTase, indicated by K_d_ values for HC (16.0 ± 2.3 μM) and CAPE (134.6 ± 14.0 μM). (Fig.6B-C).

### 3.7 Tryptophan fluorescence spectroscopy-based HC and CAPE interaction with KFDV NS5 MTase protein

The interactions of compound HC and CAPE with KFDV NS5 MTase protein were investigated using tryptophan fluorescence spectroscopy (TFS). The Synergy H1 multimode plate reader (Agilent BioTek) was utilised to monitor the intrinsic fluorescence of the KFDV NS5 MTase protein, and a progressive quenching of fluorescence intensity was observed with increasing concentrations of HC and CAPE. In TFS, a red shift in emission maxima reflects reduced hydrophobicity and increased polarity around tryptophan residues, whereas a blue shift indicates increased hydrophobicity and decreased polarity. The protein displayed a dose-dependent red shift upon binding to HC and CAPE. For KFDV NS5 MTase, K_d_ values were determined as 1.55±0.39µM for HC and 237.5±55.86 µM for CAPE. (Fig.7)

**Fig.7.**
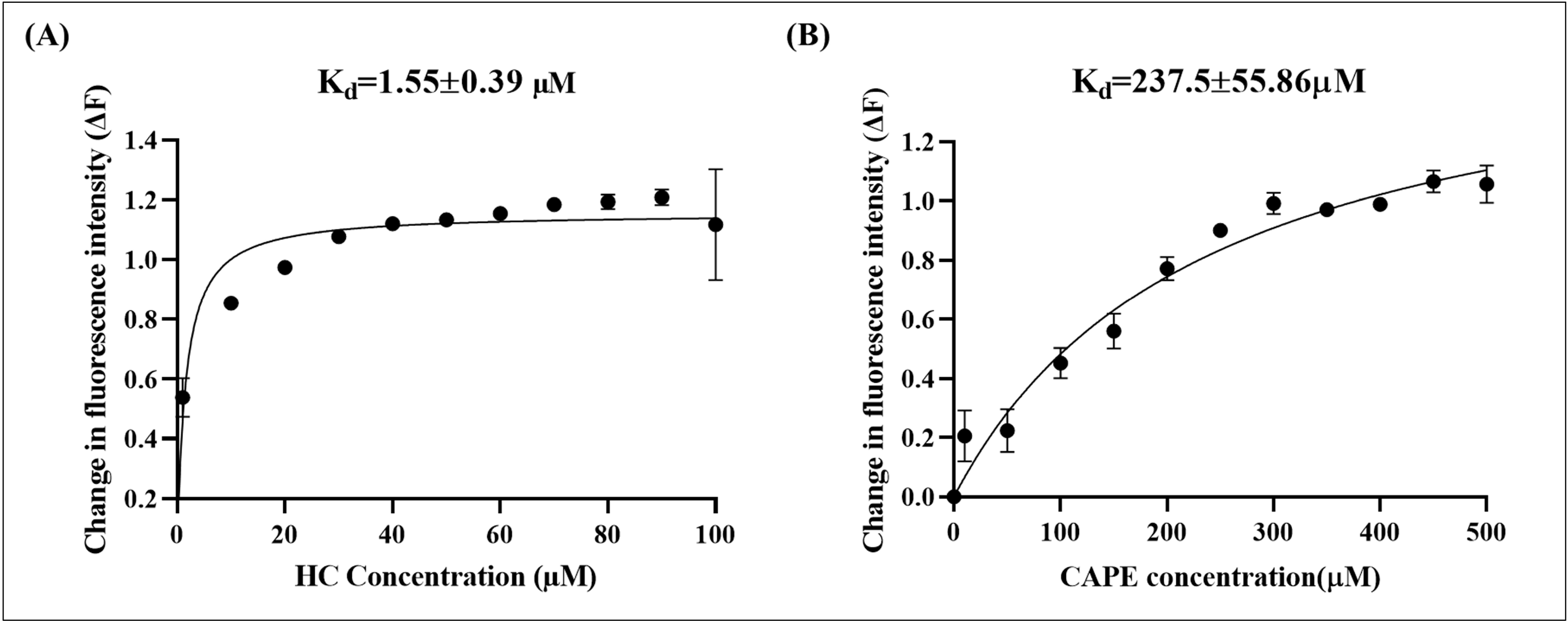
Tryptophan fluorescence spectroscopy. response indicating inhibitory effects of increasing concentrations of (a) HC, (b) CAPE with KFDV NS5 MTase protein. The standard deviation (SD) is represented as error bars derived from the means of triplicates.

## 4. Discussion

Kyasanur Forest disease virus (KFDV), an emerging tick-borne Orthoflavivirus associated with severe febrile and neurological disease. The NS5 MTase protein is a highly conserved and functionally indispensable component of the Orthoflavivirus capping machinery which is responsible for sequential catalytic activity of both N7- and 2′O-methylation of the nascent viral RNA capping and for immune evasion. The conservation of the MTase active site across Orthoflaviviruses makes it a most promising drug target for structure-guided inhibitor discovery and supports its relevance as a pan-Orthoflavivirus target.

In this study, the high-resolution structural characterisation of the KFDV NS5 methyltransferase (MTase) domain has been presented in complexes with S-adenosyl-L-homocysteine (SAH) and guanosine-5’-triphosphate (GTP), revealing canonical Orthoflavivirus MTase architecture while elucidating key interactions significant for cap methylation and substrate recognition. The KFDV MTase adopts the expected Rossmann-like fold typical of Orthoflavivirus MTases, comprising a central β-sheet flanked by α-helices and deep cleft that forms SAM or SAH-binding pocket. The overall topology closely resembles that of other Orthoflavivirus NS5 MTases, supporting the conservation of fold and mechanism across the family.

The SAH-bound structure represent state of product methyltransferase reaction and maps a dense network of hydrogen bonds, hydrophobic contacts, and ordered waters that stabilize the ligand. Key polar interactions define a well-ordered binding geometry that resembles canonical SAM recognition motifs observed in other orthoflavivirus NS5 MTases. The identification of multiple glycine and small-residue contacts (Gly60, Gly63, Gly83, Gly87, Gly88) and involvement of aromatic/hydrophobic residues (Trp89, Ile107, Ile149, Met132, Tyr135) emphasizes a binding pocket that balances precise polar contacts for positioning the methyl donor or acceptor and hydrophobic interactions for adenine stabilization.

Functionally, the observed SAH interactions support a classical methyl transfer mechanism in which the Rossmann fold positions the co-factor for methyl donation, while the local hydrogen bond network and van der Waals’ contacts likely orient the methyl group and stabilize the post-transfer product. The catalytic K-D-K-E tetrad (Lys63, Asp148, Lys185, Glu221) is present in the expected motifs and forms a positively polarized environment appropriate for stabilizing the transition state during methyl transfer. The placement of this motif confirms conservation of the catalytic core and suggests that KFDV MTase shares the same catalytic strategy as characterized for other orthoflaviviruses. The GTP-bound complex provides structural evidence for how the MTase recognizes the 5′-terminal guanosine of the cap structure. GTP occupies the RNA-binding face opposite the SAM pocket, consistent with a two-site arrangement that simultaneously accommodates the co-factor and RNA. The guanine base is stabilized by π-π stacking with Phe26, while multiple polar contacts anchor both the nucleobase and the sugar-phosphate moiety. Salt-bridge interactions from Arg30 and Arg216 to phosphate oxygens highlight the importance of basic residues for stabilizing the negatively charged triphosphate, consistent with the observed basic groove that engages RNA phosphate backbones in Orthoflavivirus MTases.

The observed contacts suggest a recognition mode optimized for the GpppA cap: stacking of the guanine ring and arginine-mediated phosphate stabilization position the 5′ guanosine for subsequent events of N7 and 2′-O methylation. The presence of SAH in the same crystal further supports a catalytically relevant arrangement in which co-factor or the product and cap substrate are accommodated simultaneously, supporting a sequential or concerted mechanism for cap methylation. The structural studies have significant implications for inhibitor design and antiviral strategies, as the detailed map of SAH and GTP contacts provides a structural framework for structure-based rational inhibitor design. Compounds that mimic the adenine or ribose-phosphate geometry, occupy the SAM-binding cleft, or bridge both SAM and cap pockets could achieve potent inhibition by blocking methyl transfer or stabilizing inactive conformations.

The two natural compounds HC and CAPE emerge as rational and mechanistically attractive lead compounds for KFDV NS5 MTase inhibition because both compounds have previously documented antiviral activity against viral methyltransferases and orthoflavivirus/alphavirus replication. In the present study, the combined four orthogonal approaches-tryptophan fluorescence spectroscopy (TFS), Isothermal titration calorimetry (ITC), fluorescence polarization (FP), and capillary electrophoresis (CE) assay provide convergent evidence that HC and CAPE directly interact with KFDV NS5 MTase and impair enzymatic activity and suppress its catalytic output. The fluorescence quenching observed in tryptophan fluorescence is consistent with ligand-induced perturbation of the local protein environment, whereas ITC establishes direct binding through thermodynamically favourable interactions. In parallel, FP assay indicates that HC and CAPE impaired the association of KFDV NS5 MTase with RNA probe, and capillary electrophoresis revealed suppression of SAH formation that indicates impairment of catalytic activity. Overall, our results provide a conserved and pharmacologically vulnerable orthoflavivirus MTase landscape that supports KFDV NS5 MTase as a high-value antiviral target and HC or CAPE as promising drug target points for next-generation pan-orthoflavivirus inhibitor design.

## 5. Conclusion

In conclusion, the present work provides structural and biophysical evidence that KFDV NS5 MTase is a conserved and druggable antiviral target, and HC and CAPE can impair its function.

The KFDV NS5 MTase structures with SAH and GTP reveal a conserved Orthoflavivirus methyltransferase architecture and define detailed residue-level interactions that mediate co-factor interaction and RNA cap recognition. These structural data deepen our understanding of cap methylation in KFDV NS5 MTase, identify residues that are critical for function, and provide a strong structural foundation for biochemical characterization and inhibitor development targeting the MTase catalytic machinery. In addition, the combined results of TFS, FP, ITC, and CE demonstrate that HC and CAPE directly interact with KFDV MTase and impair its catalytic activity, supporting their potential as broad-spectrum orthoflavivirus inhibitor lead compounds.

## 6. Acknowledgements

The authors gratefully acknowledge Department of Biosciences and Bioengineering at the Indian Institute of Technology Roorkee for access to its central facilities. They also thank the Ashok Soota Molecular Medicine Facility at IIT Roorkee and the Macromolecular Crystallographic Unit Facility (MCU) for crystallographic support. ST and PK acknowledge the Department of Biotechnology, Government of India, through the National Network Project of the Department of Biotechnology at the Indian Institute of Technology Roorkee (Project No. BT/PR40142/BTIS/137/72/2023). The authors also acknowledge financial support from the University Grants Commission (UGC), Council of Scientific and Industrial Research (CSIR), Indian Council of Medical Research (ICMR), and Ministry of Human Resource Development (MHRD). PK and ST acknowledged funding and project support from the Ministry of Education’s (MoE) Scheme for Transformational and Advanced Research in Sciences (STARS) (under project reference no. STARS2/2023-0209). The authors acknowledge and thank Bioinformatics Center (BIC) supported by Department of Biotechnology, Govt. of India (reference number: BT/PR40141/BTIS/137/16/2021).

## Notes

### Competing Interest Statement

The authors have declared no competing interest.

