## Supplementary Figure 1 and Supplementary Figure 2 for "Structural Insights and Inhibitor Discovery for Kyasanur Forest Disease Virus NS5 Methyltransferase"

***Shailly Tomar**

Professor,

Department of Biosciences and Bioengineering,

Indian Institute of Technology Roorkee,

Uttarakhand (247667), India

ORCID ID: 0000-0002-1730-003X

**Supplementary Figure 1: SDS-PAGE image of KFDV NS5 MTase protein**

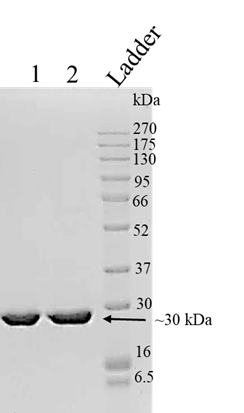

**Supplementary Figure 1: 12% SDS-PAGE analysis of purified KFDV NS5 MTase Protein fractions. Lane 1 and 2 (Ni-NTA elution fractions) ~30 kDa, and Lane 3 (protein molecular weight marker).**

**Supplementary Figure 2: Size exclusion chromatography image of KFDV NS5 MTase protein**

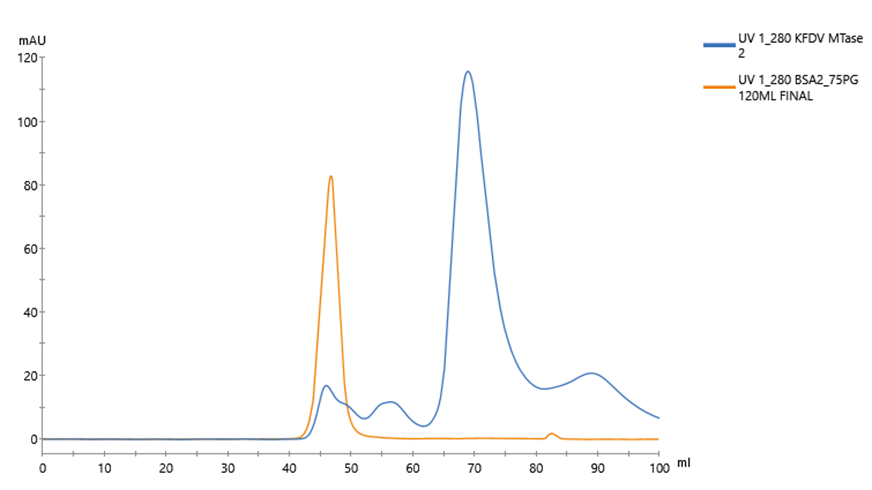

**KFDV NS5 MTase ~30 kDa**

**Peak at 68.89ml**

**BSA**

**~66.5kDa**

**Supplementary Figure 2: Size-exclusion chromatography profile on Superdex 75PG 16/600 GL column (GE Healthcare). The major peak eluting at ~68.89 mL corresponds to monomeric KFDV NS5 MTase (~MW 30 kDa), in reference to the BSA peak eluted at 46.54mL, which corresponds to 66.5kDa (monomer).**
